# Does human P-glycoprotein efflux involve transmembrane alpha helix breakage?

**DOI:** 10.1101/692988

**Authors:** Cátia A. Bonito, Maria-José U. Ferreira, Ricardo J. Ferreira, Daniel J. V. A. dos Santos

## Abstract

The occluded conformation suggested in a recent article that revealed a new inward-facing conformation for the human P-glycoprotein may not represent the closing of a gate region but instead an artifact derived from lateral compression in a too small sized nanodisc, used to stabilize the transmembrane domains of the transporter.

---

A research article published by Locher and co-workers [1] turns an important page in studying ABC transporters because it is the first cryo-EM inward-facing structure of human P-glycoprotein (P-gp), obtained with both substrate and inhibitor bound, that together with the outward-facing ATP-bound human P-gp published in 2018 [2] allow a more complete overview on the conformational changes associated to the efflux cycle. However, the claiming by the authors that authors state that “…an occluded conformation (…) in a central cavity formed by closing of a gate region consisting on TM4 and TM10” must be thoroughly discussed. Although correct from the commonly accepted mechanistic point of view, the presence of such discontinuities in both helices is strikingly different from all other published structures [3–8], in which both TM4 and/or TM10 are depicted as full helical domains. As under normal bending forces the breakage of hydrogen bonds that hold the helical structure together should not occur [9], it is important to try clarifying the underlying reason.

This specific feature had already been reported for a murine P-gp structure [8] in the presence of cyclopeptides from the QZ series. Together with ATPase and Calcein-AM transport assays, Szewczyk and co-workers proposed that the structural kink observed in TM4 (but not TM10) were induced by ligands that function more as activators of ATPase (QZ-Ala and QZ-Val) while non-activators (QZ-Leu and QZ-Phe) failed to induce the TM4 kink. However, a previous P-gp structure also co-crystallized with QZ-Val failed to show any kink on either TM4 or TM10 [3,10]. This way, the comparison between both murine and human structures suggests three additional questions: 1) if related to ligand entry, why did not TM10 kinked as TM4 after the passage of the QZ activators through the TM4/TM6 portal in the murine structure?, 2) by which gate taxol entered into hP-gp, because both TM4 and TM10 are kinked and 3) if related to ATPase activators, why did zosuquidar (inhibitor, IC_50_ 60 nm) [11] induced a similar TM4/TM10 kink as substrates do? Most intriguingly, the proposed occluded conformation also impairs the regulation of the inherent substrate specificity of P-gp by a small linker region (missing in all structures) [12,13] because it becomes unable to directly interact with substrates.

To answer all these questions, we started by comparing all P-gp (ABCB1) structures available in the Protein Data Bank (**Table 1**). It is possible to verify that, while NBD-NBD distances in murine P-gp structures are within 48 Å (4M1M) up to 65 Å (4Q9K), in the inward-facing human P-gp, this distance reduces to only 34 Å (bound to zosuquidar) or 37 Å (bound to taxol), only 7-8 Å greater than those registered for the outward-facing P-gp structures reported to date (28 Å) [2,14]. However, as these structures were obtained without the presence of ATP, how can we explain such reduction in NBD-NBD distances? More interestingly, another murine P-gp structure also obtained by cryo-EM reported a distance of 55 Å, well within the ones reported previously in the other crystallographic structures. When reporting only to the cryo-EM structures, both inward murine (PDB ID: 6GDI) and outward human (PDB ID: 6C0V) were found to have both TM4/TM10 modelled as straight helices. Furthermore, while Thongin and co-workers identified electron densities compatible with “detergent head-groups from the annular detergent micelle (…) close to two regions predicted to delineate two pseudo-symmetry-related drug-binding sites” but no TM4/TM10 kink [5], other authors reported for the outward-facing human structure that the “continuity of these helices is important to completely close the intracellular gate upon NBD dimerization, avoiding potential leakage in the outward-facing state” [2]. So, what could be the underlying reason for such a difference?

**Table 1.** Comparison of P-glycoprotein structures available in the Protein Data Bank.

| PDB ID | Organism | Ligand | Class | Conformation | NBD-NBD (Å) | Distorted TM's | Type |
| --- | --- | --- | --- | --- | --- | --- | --- |
| 3WME | *C. merolae* | -- | -- | inward | 42.59 | -- | crystal |
| 3WMG | *C. merolae* | aCAP | -- | inward | 42.59 | 4 | crystal |
| 4F4C | *C. elegans* | undecyl b-D-thiomaltopyranoside | detergent | inward | 58.82 | 3, 9, 10, 12 | crystal |
| 4M1M | *M. musculus* | -- | -- | inward | 48.03 | 12 | crystal |
| 4M2S | *M. musculus* | QZ59-Val (RRR) | inhibitor | inward | 48.32 | 12 | crystal |
| 4Q9H | *M. musculus* | -- | -- | inward | 61.85 | -- | crystal |
| 4Q9I | *M. musculus* | QZ-Ala (x2) | substrate | inward | 59.80 | 4 | crystal |
| 4Q9J | *M. musculus* | QZ-Val (x2) | substrate | inward | 62.56 | 4 | crystal |
| 4Q9K | *M. musculus* | QZ-Leu | inhibitor | inward | 65.38 | -- | crystal |
| 4Q9L | *M. musculus* | QZ-Phe | inhibitor | inward | 60.00 | -- | crystal |
| 4XWK | *M. musculus* | BDE-100 | pollutant | inward | 60.69 | -- | crystal |
| 5KPI | *M. musculus* | -- | -- | inward | 47.99 | 12 | crystal |
| 6A6M | *C. merolae* | -- | -- | outward | 27.65 | -- | crystal |
| 6A6N | *C. merolae* | -- | -- | inward | 43.19 | 4 | crystal |
| 6C0V | *H. sapiens* | ATP | nucleotide | outward | 28.28 | -- | cryo-EM |
| 6FN1 | chimera | zosuquidar (x2) | inhibitor | inward | 34.14 | 4, 10, 12 | cryo-EM * |
| 6QEX | *H. sapiens* | taxol | substrate | inward | 36.72 | 4, 10, 12 | cryo-EM * |
| 6GDI | *M. musculus* | -- | -- | inward | 55.19 | -- | cryo-EM |

\* Obtained inserted in a lipid nanodisc.

Quite remarkably, the human P-gp structures reported by Locher and co-workers are the first ones in which a nanodisc was employed to maintain a hydrophobic environment around the transmembrane helices, while in all others detergent micelles were used. Nanodiscs are derived from apolipoprotein A1, also called membrane scaffolding protein (MSP), an amphipathic protein in which the hydrophobic helices interact with the acyl chains of the lipids while the hydrophilic sides are exposed to the solvent, thus allowing the formation of a lipid bilayer patch in solution. However, it was recently reported that the nanodisc lipid internal dynamics and thermotropism may be substantially altered, abolishing gel-fluid phase transitions and increasing site-specific order parameters that can only be fine-tuned with the addition of cholesterol [15]. As it is known that P-gp reshapes the surrounding lipid environment up to a radius of 15-20 nm [16], and as the inferred nanodisc size is ~10 nm (containing 120-160 lipids) [15], under the temperature conditions required by cryo-EM and using relatively small nanodiscs, the protein conformational space may be perturbed.

Although other papers already reported murine [17,18] and human [19] P-gp activities reconstituted in nanodiscs, we identified at least two main differences in nanodisc preparation. While in biochemical experiments DMPC and *E. coli* total lipid extract was used, a mixture of brain polar lipids and cholesterol was used in the cryo-EM. More important, while in the first cases the purified P-gp, MSP and lipid ratio combination were 1:50:1750 and 1:10:800 respectively, in the cryo-EM the ratio was 1:10:350. A recent paper reporting allosteric modulation of P-glycoprotein also reconstituted the transporter in nanodiscs, also used much higher ratios, 1100:10:1 for lipid, MSP and protein respectively [20]. Therefore, the ratio used in cryo-EM seems very low and, since it also includes cholesterol, it is conceivable that the number of lipids within the cryo-EM nanodisc was smaller, which could have a direct influence on the protein architecture.

One could argue that human P-gp is intrinsically different from murine P-gp (therefore the necessity of using nanodiscs) and that this was not the only protein obtained in a nanodisc. Searching through the most recent literature, we find that a V-ATPase Vo proton channel (protein:MSP:lipid ratio 1:50:1250, PDB ID: 6C6L), a voltage-activated Kv1.2-2.1 paddle chimera channel (ratio 1:10:400, PDB ID: 6EBK) or a human α1β3γ2L GABAA receptor (MSP:lipid ratio ~1:100, PDB ID: 6I53). However, and besides the higher MSP:lipid ratio that is always used in the assembly of the nanodiscs, an important difference is that all these examples possess a radial symmetry for the portion embedded in the membrane while P-gp has a pseudo two-fold symmetry. Quite interestingly, both TM4 and TM10 are located across the smaller symmetry axis, which also indirectly supports the assumption that a highly restrained environment as in cryo-EM conditions may induce such structural deformations.

To provide a proof of concept, we have generated a series of molecular dynamics (MD) simulations to mimic the effect of lateral compression by a nanodisc under low-temperature conditions. To speed up the calculations all waters were removed, phosphate atoms were spatially restrained in *z* (xy corresponds to the membrane plane) to prevent membrane disassembling and a lateral pressure of 1.5 bar *(xy* only) was applied, using an anisotropic pressure coupling scheme. To further promote lipid order parameters, we additionally decreased the membrane temperature (280K) while keeping the protein at 310K. Two systems were simulated, in the absence or presence of a ligand. From **Figure 1**, it is possible to verify that, in the absence of ligands, only one helix was found to bend, in this case TM10 (**Figure 1A**), similar to what was found for the murine structures published by Szewczyk and co-workers. When in the presence of a large ligand as cyclosporin (with a MW above 600 Da, as taxol), both TM4 (**Figure 1B**) and TM10 (**Figure 1C**) helices were found to bend inwards, immediately below the ligand and apparently ‘closing’ the access to additional molecules. This is clear from the decrease in the root-mean square deviation between both helices regarding the human P-gp cryo-EM structure and the human P-gp homology model, shifting from 4.61 Å to 3.24 Å in TM10 and from 4.21 Å to 2.89 Å in TM4. The distance between TM4 and TM10 was also found to decrease during the 100 ns run, from 18.8 Å to 13.4 Å. Thus, our simulations suggest that by applying a mild increment on the lateral pressure, TM4 and TM10 tend to bend inwards immediately below the substrate, as observed in the cryo-EM structure by Locher and co-workers.

**Figure 1.**
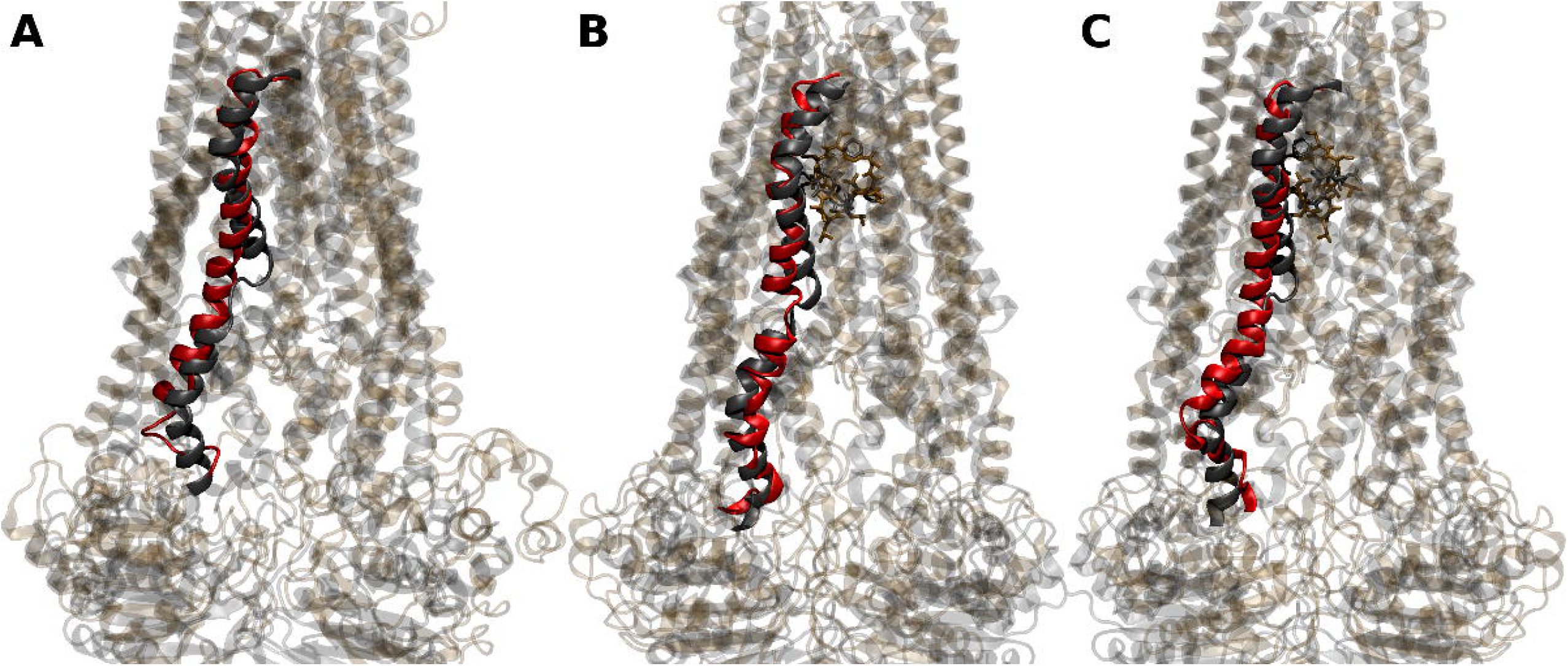
Superimposition of human P-gp (PDB ID: 6QEX, grey) with the final conformations of molecular dynamics runs starting with a homology P-gp model (red). (A), *apo* structures (TM10 depicted); *holo* structures depicting taxol (grey) and cyclosporine (ochre) together with TM4 (B) or TM10 (C).

Interestingly, in the murine structures previously reported only when two QZ molecules were found at the drug-binding site (combined MW, 1212 Da) a TM4 kink was present, and in our simulations only in the presence of bound molecules such kinks were formed. More surprisingly, it was only necessary to apply a slightly higher pressure (1.1 bar) to initiate such changes, becoming more apparent when using 1.5 bar. Furthermore, we additionally observed that this effect seems to occur quite fast in the time scale of the simulations, as we were able to obtain similar results using short and harsher simulation conditions (1 ns each) or longer simulations (100 ns), using more mild conditions (0.2 bar increments over 0.5 ns until reaching 1.5 bar). In other words, even when the pressure increment is slower and occurs over a longer time, the distortion on TM4 and 10 occurs promptly when in the presence of a large ligand at the internal drug-binding pocket.

Therefore, it is our opinion that, and unlike the initially postulated hypothesis, the observed ‘gate closing’ in the recently published cryo-EM human P-glycoprotein structure may be an artifact from a too small nanodisc construct used for stabilizing the transmembrane domains of the transporter. Under certain cryo-EM conditions, the internal lipid dynamics provided by the nanodisc may render unfavorable conditions for the structural stability of the transporters’ architecture, leading to severe distortions of TM4 and TM10 helices. This also upholds the importance of a careful choice of the experimental conditions and protocol to be used to obtain reliable models of membrane proteins since these cryo-EM structures are establishing new and improved starting points for improved computer simulations [21–23].

## Acknowledgement

Fundação para a Ciência e Tecnologia (FCT) is acknowledged for financial support (PTDC/MED-QUI/30591/2017, PTDC/MED-QUI/28800/2017 and SAICTPAC/0019/2015). Cátia A. Bonito also acknowledges FCT for the Ph.D grant SFRH/BD/130750/2017.

## Author contributions

R. J. F. and D. J. V. A. S. formulated the hypothesis. C.A.B., R. J. F. and D.J.V.A.S. written the manuscript. R. J. F. performed the MD simulations and rendered the images. All authors revised the manuscript and agree with its final form.

